# Hexanucleotide repeat expansions in C9orf72 alter microglial responses and prevent a coordinated glial reaction in ALS

**DOI:** 10.1101/2022.10.26.513909

**Authors:** Pegah Masrori, Baukje Bijnens, Kristofer Davie, Suresh Kumar Poovathingal, Annet Storm, Nicole Hersmus, Laura Fumagalli, Ludo Van Den Bosch, Mark Fiers, Dietmar Rudolf Thal, Renzo Mancuso, Philip Van Damme

**Affiliations:** KU Leuven - University of Leuven, Department of Neurosciences, Leuven Brain institute (LBI), Leuven, Belgium; Laboratory of Neurobiology, VIB Center for Brain & Disease Research, Leuven, Belgium; University Hospitals Leuven, Department of Neurology, Leuven, Belgium; Microglia and Inflammation in Neurological Disorders (MIND) Lab, VIB Center for Molecular Neurology, VIB, Antwerp, Belgium; Department of Biomedical Sciences, University of Antwerp, Antwerp, Belgium; Single Cell Bioinformatics Expertise Unit (CBD), VIB Center for Brain & Disease Research, Leuven, Belgium; Single Cell Analytics & Microfluidics Core, VIB Center for Brain & Disease Research, Leuven, Belgium; VIB Center for Brain & Disease Research, Leuven, Belgium; Laboratory for Neuropathology, Department of Imaging and Pathology, Leuven Brain Institute (LBI) KU Leuven, Leuven, Belgium; Department of Pathology, University Hospitals Leuven, Leuven, Belgium

## Abstract

Neuroinflammation is an important hallmark in amyotrophic lateral sclerosis (ALS). Experimental evidence has highlighted a role of microglia in the modulation of motor neuron degeneration. However, the exact contribution of microglia to both sporadic and genetic forms of ALS is still unclear. We generated single nuclei profiles of spinal cord and motor cortex from sporadic and *C9orf72* ALS patients, as well as controls. We particularly focused on the transcriptomic responses of both microglia and astrocytes. We confirmed that *C9orf72* is highly expressed in microglia and shows a diminished expression in carriers of the hexanucleotide repeat expansion (HRE). This resulted in an impaired response to disease, with specific deficits in phagocytic and lysosomal transcriptional pathways. Astrocytes also displayed a dysregulated response in *C9orf72* ALS patients, remaining in a homeostatic state. This suggests that C9orf72 HRE alters a coordinated glial response, which ultimately would increase the risk for developing ALS. Our results indicate that *C9orf72* HRE results in a selective microglial loss-of-function, likely impairing microglial-astrocyte communication and preventing a global glial response. This is relevant as it indicates that sporadic and familial forms of ALS may present a different cellular substrate, which is of great importance for patient stratification and treatment.

## Introduction

Amyotrophic lateral sclerosis (ALS) is a neurodegenerative disease characterized by progressive motor neuron loss (MNs) in the motor cortex, brainstem, and spinal cord (1). Motor neuron loss is accompanied by neuroinflammation, and changes in both microglia and astrocytes (2). ALS typically presents with focal muscle weakness, which spreads to neighboring body regions. Extramotor manifestations, such as cognitive and behavioral impairment, occur in a subset of patients. Effective therapies are lacking, and the median survival after symptom onset is limited to 3 years (1). The widespread nature of this disease, with contribution from distant central nervous system (CNS) regions, represents a challenge when trying to elucidate the underlying cellular mechanisms. To achieve a deep understanding of ALS pathophysiology, it is essential to investigate cell-specific disease mechanisms acting both in the spinal cord and the motor cortex.

ALS has a strong genetic component, where the most common monogenic cause is the GGGGCC repeat expansions in the *chromosome 9 open reading frame 72* gene (*C9orf72*) (3), with a repeat length ranging from 38 to approximately 1600 repeats (4, 5). *C9orf72* presents with multiple transcript variants that are differentially enriched across cell types in the brain (6, 7). Both loss-of-function (LOF) and toxic gain-of-function (GOF) mechanisms in neuronal and non-neuronal cells have been implicated in the disease pathogenesis (8). *C9orf72* is predominantly expressed in myeloid cells, including microglia (6). Whereas the exact mechanism by which the *C9orf72* mutations alters microglia function is not fully understood (9), experimental studies have shown that its deletion leads to altered immune responses, exacerbated myeloid proliferation and lysosomal dysfunctions (10, 11).

In this study, we provide an unparalleled single-nuclei RNA sequencing resource from paired motor cortex and spinal cord samples from sporadic ALS without any known gene mutation (sALS), ALS patients with a *C9orf72* mutation (C9-ALS) and controls. We show that mutations in *C9orf72* prevent the transition of microglia towards reactive cell states, and block a coordinated multi-cellular response to disease, when compared to sALS. While sALS and C9-ALS are clinically indistinguishable, our findings suggest that cellular substrate underlying disease might be fundamentally different. Also, our work allows interrogating the contribution of cellular subtypes involved in disease pathogenesis and cell-specific dysregulated pathways. We believe this unique resource and approach will lead to the identification of new neuroinflammatory biomarkers, better patient stratification and novel, more specific, disease-modifying strategies to tackle ALS.

## Results

We performed single nuclei RNA (snRNA)-sequencing on motor cortex and spinal cord samples from 5 sALS, 5 C9-ALS and 5 controls. The experimental setup is illustrated in Figure 1a. The demographic data of participants are shown in Supplementary Table 1. To minimize potential transcriptomic artefacts (12), we only used samples with a mean post-mortem interval of up to 9 hours.

**Figure 1.**
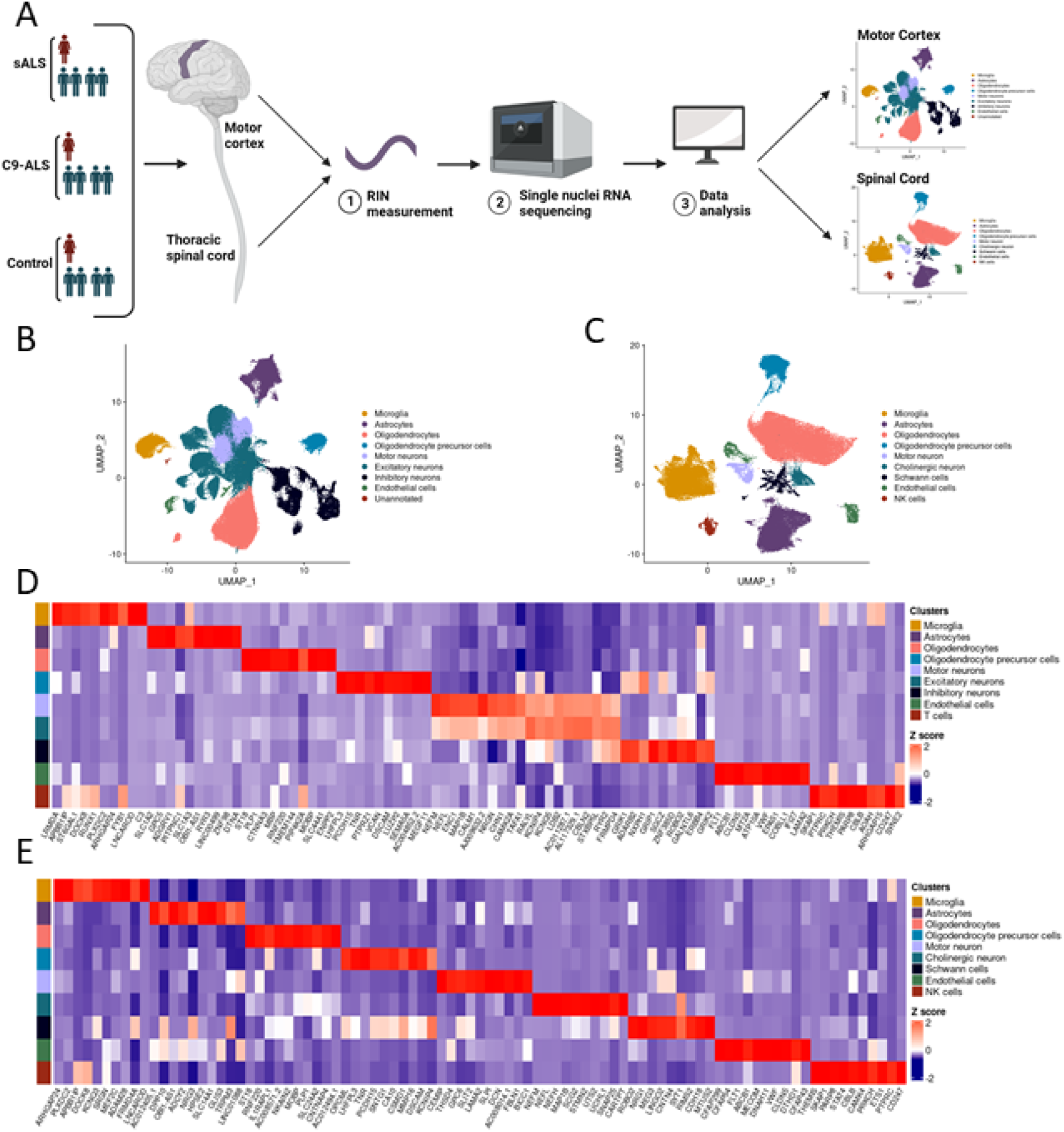
sn-RNA sequencing of human post mortem motor cortex and spinal cord of ALS patients. (**a**) experimental outline of this study. (**b**) UMAP plot visualizing 93,176 nuclei isolated from post-mortem cortical samples of ALS patients (C9-ALS and sALS) and controls. Cells are colored according to their assigned cluster identity: microglia, astrocytes, oligodendrocytes, oligodendrocyte precursor cells, motor neurons, excitatory neurons, inhibitory neurons, endothelial cells and T-cells. (**c**) UMAP plot visualizing 63,076 nuclei isolated from post-mortem spinal cord samples of ALS patients (C9-ALS and sALS) and controls. Cells are colored according to their assigned cluster identity: microglia, astrocytes, oligodendrocytes, oligodendrocyte precursor cells, motor neurons, cholinergic neurons, Schwann cells, endothelial cells and an unannotated cluster. Heatmap showing the top 10 most expressed genes per cluster of the (**d**) motor cortex and (**e**) the spinal cord.

### Optimization of single nuclei isolation and sequencing from post-mortem human samples

Recent advances in the isolation and sequencing of snRNA have demonstrated that, with optimal buffer/detergent compositions, the assay performance and gene detection sensitivity can be significantly enhanced. We first evaluated three different strategies for single nuclei isolation: the protocol by Habib et al. (13), Tween with salts and Tris (TST) (14), and Soma-seq (15) (Methods Online). After nuclei isolation, we performed snRNA sequencing of spinal cord and motor cortex from sporadic and C9orf72 ALS patients, as well as controls and compared the three methods for basic parameters including number of reads, genes and UMIs. Based on this analysis we concluded that the Soma seq method shows the best performance (Supplementary Figure 1c), and therefore we used this method for all the following experiments.

### Cellular diversity in the spinal cord and motor cortex

We investigated the cellular diversity across the motor cortex spinal cord. We analyzed 156,252 nuclei, after removal of low-quality nuclei and doublets. The full dataset from both pre- and post-batch correction is shown in Supplementary Figure 1.

In the spinal cord, we identified 9 major transcriptionally distinct nuclei clusters accounting for all the cell types present in the spinal cord (Figures 1c, e). We annotated these clusters into: microglia (*ARHGAP24, PLXDC2, KCNQ3, SRGN*), oligodendrocytes (*ST18, IL1RAPL1, MOBP, SLC24A2*), astrocytes (*NRG3, DPP10, HPSE2, SCL14A1*), oligodendrocytes precursor cells (OPCs, *OPCML, TNR*,, *MMP16*), Schwann cells (*ROBO2, NRG1, MEG3, CNTN4*,), motor neurons (*CEMIP, SLT2, LAMA2, DCN*), cholinergic neurons (*NEFM, MAP1B, STMN2, SNAP25*), endothelial cells (*FLT1, ABCB1, CLDN5, CFAP43*), and Natural Killer cells (*THEMIS, SKAP1, PARP8, STAT4*).

On the other hand, we identified 9 major transcriptionally-distinct nuclei clusters in the motor cortex (Figures 1b, d), including oligodendrocytes (*ST18, PLP1, MBP, MOBP*), astrocytes (*SLC1A2, SLC1A3, RYR3, DTNA*), excitatory neurons (*RALYL, CBLN2, KCNQ5, LDB2*), microglia (*LRMDA, ST6GAL1, RUNX1, C3*), motor neurons (*NEFM, NEFL, ENC1, MAP1B*), OPCs (*LHFPL3, TNR, VCAN, DSCAM*), inhibitory neurons (*MEGF11, GRIP1, ROBO2, GLANTL6*), endothelial cells (*ABCB1, CLDN5, MT2A, IFI27*) and T cell (*SKAP1, PTPRC, CD247, SYNE2*).

We did not observe any significant differences in the distribution of patients and experimental conditions across clusters (Supplementary Figures. 1d, e).

### The C9orf72 hexanucleotide expansion diminishes microglial activation

We subset, integrated and re-clustered the microglial nuclei from the spinal cord (n=13,980) and motor cortex (n=4,845 cells) (Supplementary Table 2). After removing peripheral and other CNS macrophages, as well as proliferating cells, (Figures 2a, b, and Supplementary Figure 2). The microglia subclusters were all characterized by a distinct transcriptomic signature (Figures 2c). Cluster MG1 closely resembled homeostatic microglia (*CX3CR1, P2RY12*) (16), and presented lower levels of known activation markers (*C1, IL1B, APOE, CTSB*) (17, 18). Cluster MG2 appeared to be a transitioning state that retained expression of homeostatic markers a more intermediate state, with expression of homeostatic markers (*CX3CR1, P2RY12*) and intermediate levels of state transition markers (*SPP1, CD14, IL1B, C*3). We captured a microglia subset, MG3, very closely related to disease/damage associated microglia (DAM) described in multiple disease scenarios (17, 19). This cluster was characterized by higher levels of *CD74, APOE, TREM2, C1QA, CTSD*. Cluster MG4 was consistent with previously described axon tract associated microglia (*LGAL3, GPNMB, MITF, CTSD*) (20). Cluster MG 5 displayed an expression profile related to hypoxia (*HIF1A, VEGFA, NAMPT, ELL2, CD83*), which has been previously described in brains with glioblastoma multiforme (21). Lastly, cluster MG 6 closely resembled cytokine response microglia (CRM) with key markers being *CCL2, CCL3* and *CCL4* (22). Distributions of these clusters over all three conditions are shown in Figure 2d.

**Figure 2.**
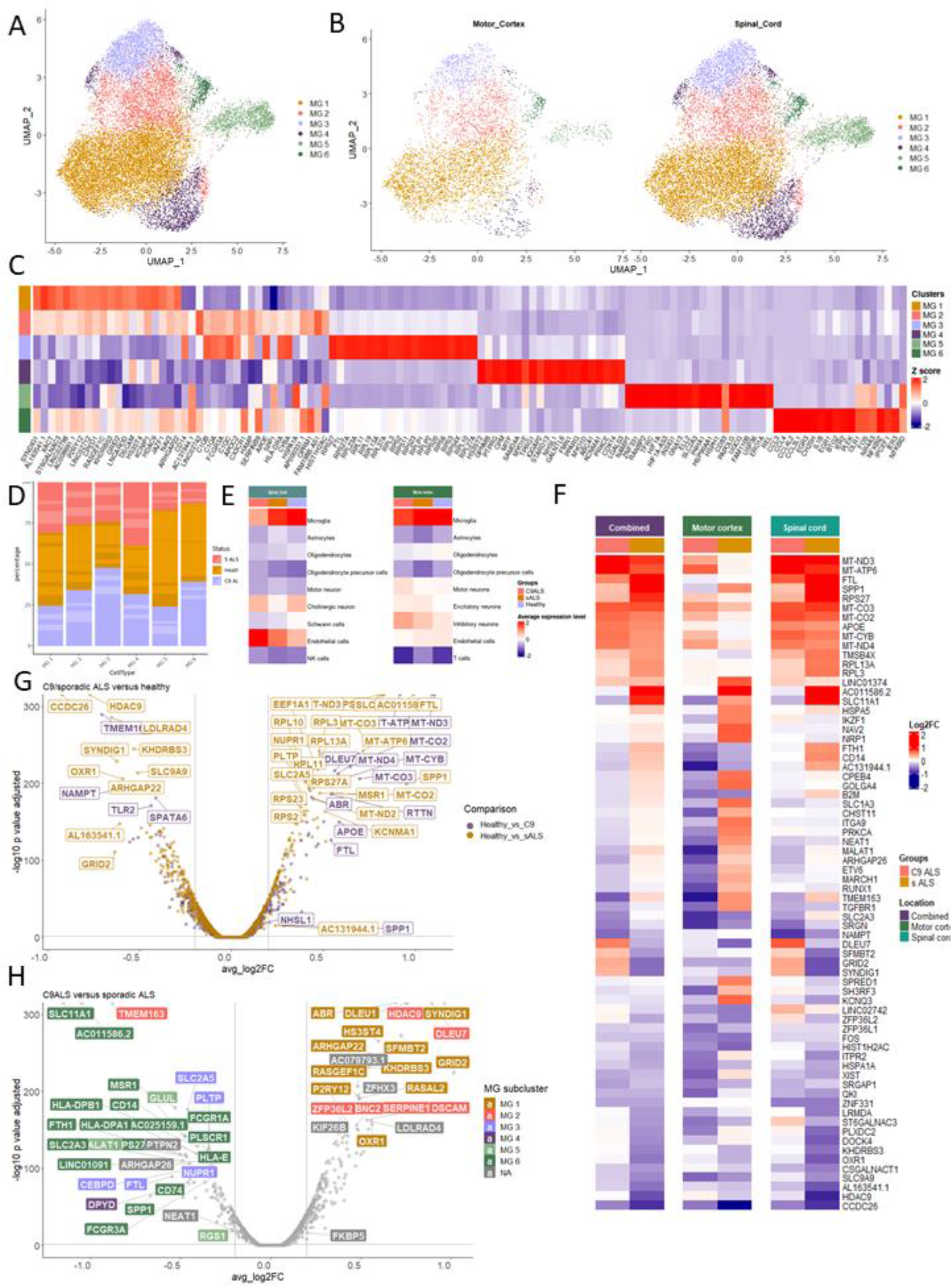
C9orf72 HRE induces changes in the microglial transcriptomic profile compared to sALS. (**a**) UMAP plot visualizing 16,763 microglia nuclei from post-mortem cortical samples of ALS patients (C9 and sALS) and controls. Microglia subsets are colored based on their assigned cluster into: MG1, MG2, MG3, MG4, MG5, MG6. (**b**) UMAP plot displaying the specific distribution of 4,216 nuclei from the motor cortex and 12,547 nuclei of the spinal cord. (c) heatmap showing the top 20 most expressed genes per cluster. (**d**) proportion of cells (y-axis) from each genotype in the different microglial subclusters, alternating shades indicate different individuals (x-axis). (**e**) heatmap depicting the Z-scored average expression of *C9orf72* in both motor cortex and spinal cord. (**f**) heatmaps showing the log2FC of differentially expressed genes (p-value adjusted <0.05) of both C9-ALS or sALS patients compared to healthy controls when comparing pooling the motor cortex and the spinal cord. (**g**) Volcano plot depicting 2 pairwise comparisons: between C9-ALS and healthy individuals, and sALS and healthy individuals for the combined microglial dataset (DESeq2 negative binomial distribution, p-values adjusted with Bonferroni correction based on the total number of genes in the dataset). Vertical lines depict Log2FC threshold of 0.2 whereas the horizontal sets the adjusted p-value threshold at < 0.05). (**h**) Volcano plot depicting the pairwise comparisons of C9-ALS vs. sALS for the combined microglial dataset (DESeq2 negative binomial distribution, p-values adjusted with Bonferroni correction based on the total number of genes in the dataset). Vertical lines depict Log2FC threshold of 0.2 and the horizontal line, the adjusted p-value threshold at < 0.05). Colors indicate marker genes of microglia clusters.

Haploinsufficiency of C9orf72 protein, as a result of the hexanucleotide repeat expansion (HRE), is a key proposed disease mechanism - along with aggregating expanded RNAs and dipeptide repeat proteins (DPRs) - in ALS (23). We wondered whether HRE in the *C9orf72* had an impact on human microglia. We assessed the levels of *C9orf72* and confirmed the HRE leads to reduced *C9orf72* expression levels (Figure 2e). Next, we performed a separate differential expression analysis between C9-ALS vs. controls, and sALS vs. controls (Figure 2f and Supplementary Table 3). We observed strong differences between experimental groups, with C9-ALS microglia displaying a much lower extent of response to ALS pathology in the motor cortex and especially the spinal cord. Comparison between C9-ALS and sALS microglia showed that while sALS microglia display transcriptomics changes consistent with a reactive state (*SPP1, CD14, HLA, FCGR1A. FCGR3A*), C9-ALS microglia remained vastly homeostatic (*P2RY12, INPP5D, BLNK, SERPINE1, SORL1* (Figure 2g, h). This suggests that the HRE in *C9orf72* impairs microglial cell state transition, consistent with a partial loss of function phenotype.

We have recently mapped the transcriptomic changes occurring in human microglia during amyloid-b pathology using a xenotransplantation mouse model (24). To shed light on whether *C9orf72* HRE modifies the microglial phenotype, we performed a gene enrichment analysis and mapped those transcriptomic signatures in our dataset). We observed the C9-ALS microglia were impaired in the transition from homeostatic to DAM and antigen-presenting HLA phenotypes (Figure 3a). This is consistent with previous observations in mouse models of C9orf72 deficiency, supporting the idea of ablation of these specific phenotypes as a result of the C9orf72 haploinsufficiency (Supplementary Figure 3) (7, 25-27). These data also indicate that there might be strong similarities between the contribution of microglia across different neurodegenerative disorders (21, 24, 28-30). To confirm our findings, we repeated our gene enrichment analysis using several specific transcriptional pathways that account for different aspects of microglial function. Whereas sALS microglia showed increased transcription of genes involved in immune, lysosomal and phagocytic pathways, C9-ALS microglia remained similar to the controls (Figures 3b, c). On the contrary, C9-ALS microglia displayed an enrichment in genes related to ribosomal function (Figures 3b, c), which is consistent with previous data reporting similar findings in *C9orf72* derived ALS astrocytes (31). Interestingly, we observed that most of the changes occurred in the spinal cord, with only mild alterations in the motor cortex (Figures 3a-c). Whereas these alterations may reflect a different extent of pathology in both regions, it could also be consequence of intrinsic regional differences in the microglia (32).

**Figure 3.**
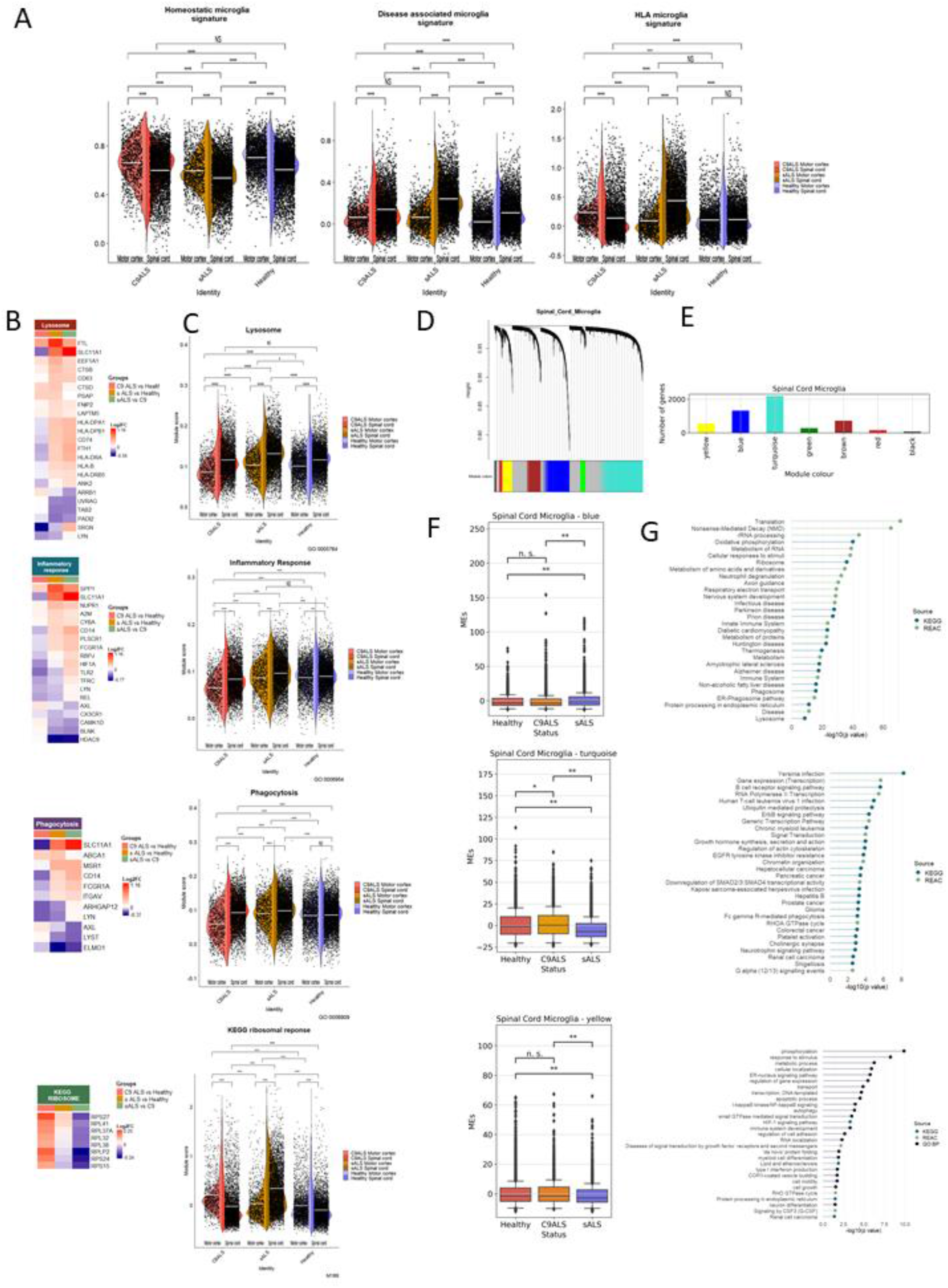
Microglia from C9orf72 HRE patients show altered functional pathways. **(a)** Seurat module scores of disease-associated (DAM), antigen-presenting HLA, and homeostatic microglia phenotypes as described by Mancuso et al., 2022. **(b)** heatmaps depicting Z-scored average expression of differentially expressed genes (log2FC < +/-0,20, p-value adjusted <0,05) of gene sets of lysosomes, inflammatory response, phagocytosis and KEGG ribosomes. **(c)**, Seurat module scores of GO terms from the lysosomes, inflammatory response, phagocytosis and KEGG ribosome. **(d)** dendrogram showing the result of unsupervised hierarchical clustering of modules identified by WGCNA. **(e)** barplots depicting the number of genes per module **(f)** boxplots visualizing the module eigengenes (y-axis) per genotype (x-axis) for the blue, turquoise and yellow cluster of the spinal cord. **(g)** pathway enrichment analysis of the WGCNA clusters blue, turquoise and yellow, with the associated GO term (y-axis) and –log10(p-value) (x-axis).

Next, we performed weighted gene co-expression network analysis (WGCNA) to confirm the alterations induced by *C9orf72* HRE in microglia. We focused on the spinal cord, where we found the most pronounced changes in our differential expression analysis. We identified 7 distinct co-expression clusters, many of them with differential significant enrichment across sALS, C9-ALS and controls (Figures 3d-f and Supplementary Figure 4). The blue cluster appeared particularly relevant, as it contained a high number of genes (Figure 3e) and was significantly enriched in sALS vs. both C9-ALS and controls (Figure 3f). Genes in this cluster were associated with functional pathways related to the immune system, phagocytosis, autophagy-lysosomal function, ubiquitin-proteasome system, oxidative stress and apoptosis confirming a strong immune component in the pathogenesis of sALS (Figure 3g). C9-ALS microglia showed no enrichment in these pathways, consistent with our findings on *C9orf72* HRE leading to an impaired microglial response (Figure 3g; Supplementary Figure 4). On the other hand, we observed an enrichment of clusters turquoise and yellow in the C9-ALS microglia compared to both sALS and controls (Figure 3f). These clusters encompass general biological pathways such as cell proliferation, migration, general metabolic processes, protein biogenesis and degradations are particularly involved, and as opposed to the blue cluster, without a predominant presence of the immune response (Figure 3g). Altogether, our data shows that the *C9orf72* HRE prevents the transition of microglia towards reactive cell states and induces perturbations in transcriptional pathways different than in sALS, therefore suggesting a different cellular mechanism underlying sporadic and inherited forms of ALS.

### Reduced microglial activation correlates with an impaired astrocytic reaction

Microglia-astrocyte interactions have been implicated in multiple neurodegenerative disorders and under inflammatory conditions (33). It is well accepted that microglia can interact and modulate the response of astrocytes (34-37), although the underlying molecular mechanism of this interplay remain unclear. As *C9orf72* is primarily expressed in the microglia (38), we hypothesize that in ALS, *C9orf72* HRE induces direct alterations in the microglia, affecting their ability to properly communicate and modulate the activation of astrocyte. To address this, we first subset, integrated and re-clustered both spinal and cortical astrocytes to define their transcriptomic landscape (Supplementary Figure 5). We observed similar heterogeneity compared to the microglia, with 8 distinct transcriptional states across the spinal cord (n=12,978) and motor cortex (n= 9,657cells) (Figures 4a-c, and Supplementary Table 2). We found a number of clusters consistent to homeostatic astrocytes, with expression of genes related to synaptic support and glutamate metabolism (AS1: *SLC1A2, NRXN1, CABLES1*) (39), interaction with neurons and axonal guidance (AS3: *TNN, NRXN3, LAMA2, PLEKHA5*) (39), or myelin formation, synaptic formation and short term plasticity (AS6: *GRID2, CNTNAP5, ERFI1, SHISA9, KCND2*) (31). Cluster AS7 was linked to axon guidance, pruning of inappropriate synapses and development of neural circuits (*GNA14, NRP1, GFRA1, CHI3L1, CPNE8*) (40). We also found several transcriptomic phenotypes consistent with astrocyte activation. Cluster AS4 showed expression of stress markers (*HES1, FOS, JUN, ID3*) (41). Cluster AS5 displayed reactive astrocytes markers (*NAMPT, CADPS, LPAR1, SERPINA3, EMP1, CHI3L2, RASGEF1B*) (39, 42). Cluster AS8 was characterized by high expression levels of reactive astrocyte markers (*SERPINA3, EMP1, CHIL3L2*) as well as multiple immune-related genes (*CLU, VIM, B2M, APOE, AQPI, C3*) (43-45). AS2 appears to be a transitioning state that retains expression of AS1 markers but also displays an enrichment of mitochondrial and ribosomal transcripts, along with intermediate levels of AS3 (*TTN, NRXN3, LAMA2, PLEKHA5*) and AS8 markers (A*POE, B2M, AQP4, SERPINA3*) (Figures 4a-c). Distributions of these clusters over all three conditions are shown in Figure 4d.

**Figure 4.**
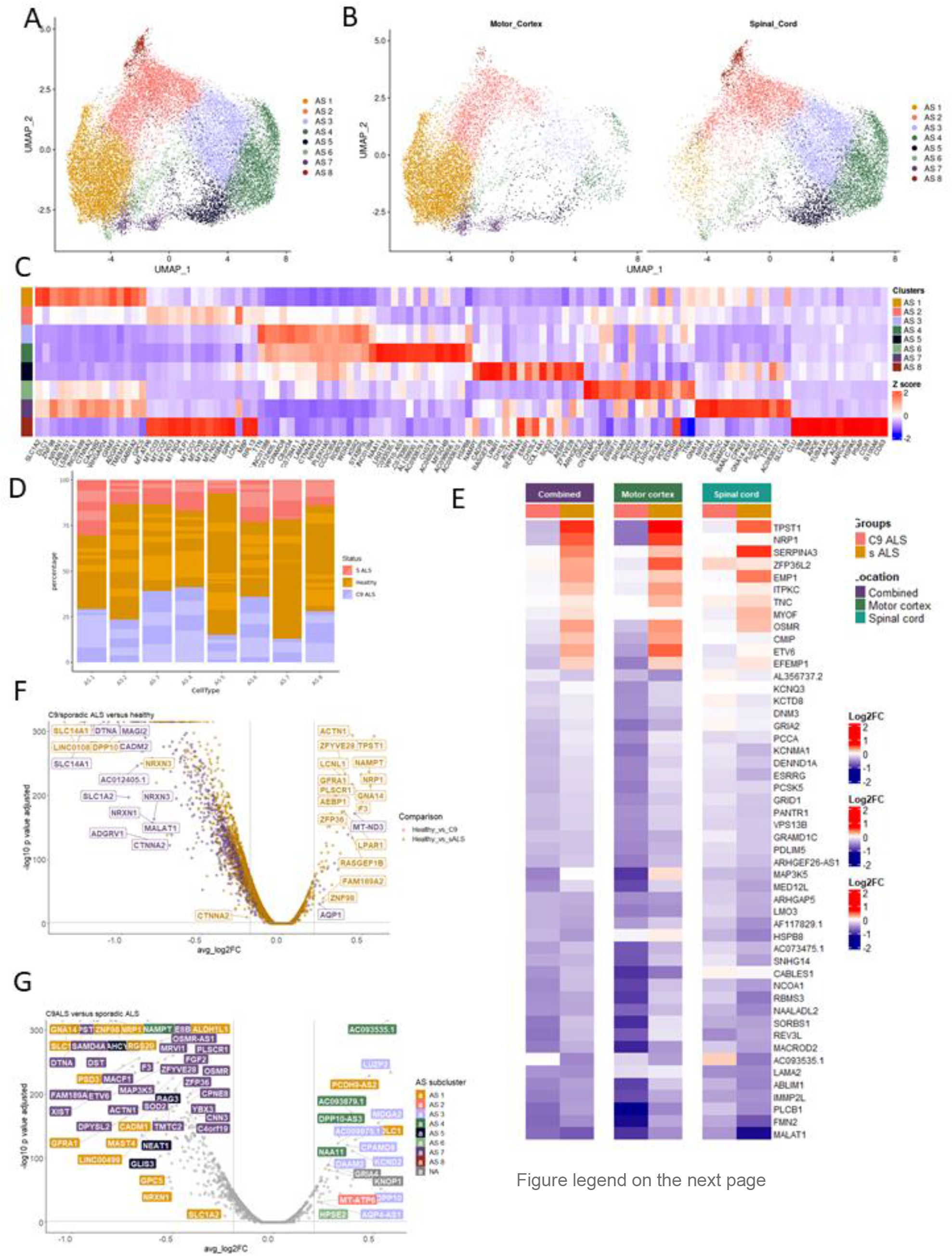
Astrocytes are differentially affected in C9-ALS compared to sALS. **(a)** UMAP plot visualizing 22,635 astrocytes from post-mortem cortical samples of ALS patients (C9-ALS and sALS) and controls. Astrocyte subsets are colored based on their assigned cluster: AS1, AS2, AS3, AS4, AS5, AS6, AS7, AS8. **(b)** UMAP plot displaying the specific distribution of the 9,657 cortical and the 12,978 spinal nuclei. **(c)** Heatmap showing the top 15 most expressed genes per cluster. **(d)** proportion of cells (y-axis) from each genotype in the different astrocyte subclusters, alternating shades indicate different individuals (x-axis). **(e)**, heatmaps showing the log2FC of differentially expressed genes (p-value adjusted <0.05) of both C9-ALS or sALS compared to healthy controls. **(f)** Volcano plot depicting 2 pairwise comparisons between C9-ALS and controls, and sALS and controls for the combined astrocytes dataset (DESeq2 negative binomial distribution, p-values adjusted with Bonferroni correction based on the total number of genes in the dataset). Vertical lines depict Log2FC threshold of 0.2 and the horizontal line, the adjusted p-value threshold at <0.05). **(g)** Volcano plot showing a pairwise direct comparisons of C9-ALS vs. sALS for the combined astrocyte dataset (DESeq2 Wilcoxon Rank sumtest, p-values adjusted with Bonferroni correction based on the total number of genes in the dataset). Vertical lines depict Log2FC threshold of 0.2 and the horizontal line, the adjusted p-value threshold at < 0.05). Colors indicate marker genes of astrocyte clusters.

We assessed for potential alterations in the response of astrocytes by performing a differential expression analysis between C9-ALS vs. controls, and sALS vs. controls (Supplementary Table 4). Similarly to what we observed in microglia, we found an overall lower extent of transcriptional changes in C9-ALS vs. sALS astrocytes (Figures 4f-e) sALS astrocytes exhibited an upregulation of genes associated to a reactive state (*SERPINA3, RASGEF, LPAR1, TPST1, CHI3L, EMP1*), as opposed to C9-ALS astrocytes which more often retained a homeostatic state (*AQP1, DLC1, SLC35F1*) (Figures 4f-g). These data indicate a *C9orf72* HRE-mediated defective engagement in a coordinated response in ALS pathology indicative of an aberrant microglia-astrocyte interaction.

## Discussion

Neuroinflammation is a key pathological hallmark for ALS (9, 46). Experimental (47, 48) and genetic evidence (46, 49, 50) indicate that microglia and neuroinflammation are active players - rather than simple bystanders - in the pathogenic events that lead to motor neuron degeneration in ALS. Our study confirms that *C9orf72* is mainly expressed in microglia in the human brain (6), and that *C9orf72* HRE leads to reduced expression. We report that *C9orf72* HRE impairs microglial cell state transition towards DAM and HLA cell states and induces alterations in transcriptomic pathways related to phagocytosis and lysosomal function. We also show that astrocytes from *C9orf72* HRE carriers display a general lower extent of transcriptomic change compared to ALS, and hypothesized that this could be the result of a defective microglia-astrocyte communication that may hinder a coordinated neuroinflammatory response to the disease. These findings suggest that patients with a *C9orf72* HRE may require a personalized intervention to modulate the neuroinflammatory response.

There are several proposed mechanisms explaining the causative role of *C9orf72* in ALS (3, 5, 51), and LOF due to reduced *C9orf72* mRNA and protein levels is one of them (23, 52, 53). Several studies have reported widespread alterations in microglia (and other myeloid cells) after conditional (54-57) or complete *C9orf72* deletion (10, 26, 58, 59), including transcriptional alterations, an exacerbated production of inflammatory cytokines, lysosomal dysfunction (6, 52, 60-62). Similarly, we also find that diminished levels of C9orf72 result in an impaired transition into DAM and HLA responses, and reduced engagement in phagocytic and lysosomal transcriptional programs. This is not a unique phenotype of microglia harboring *C9orf72* mutations, and have been observed before in pathogenic genetic variants in *TREM2* and *APOE* (24). These findings indicate that the inflammatory substrate and microglial contribution is different in *C9orf72* and sALS pathology, and additional insights will be crucial to define tailored therapeutic strategies for groups of patients with different pathogenic contributors.

Recent studies have suggested a synergistic effect of *C9orf72* LOF and GOF (63). In vivo studies showed that *C9orf72* deficiency leads to exacerbation of motor phenotypes and premature death in C9orf72 GOF mouse models (64, 65). It has been proposed that reduced *C9orf72* levels might further enhance the pathological accumulation and toxicity of DPRs (66, 67) by altering autophagy (68). However, this potential synergism between LOF and GOF has been only investigated in neurons (65). Based on our observations, it is plausible that these two mechanisms act via different cell types instead, with GOF altering neuronal function, and LOF leading to a microglial defect, exacerbation of neuronal alterations and a clinical phenotype. Whereas the mechanisms underlying the cell specific impact of C9 HRE are unclear, it is plausible that different cell types use different *C9or72* transcript variants, therefore manifesting different functional consequences of carrying the pathogenic expansion. Supporting this idea, it has been reported that while neurons, oligodendroglia, and astrocytes use transcript variants containing exons 1a and 1b, microglia are enriched in transcripts including exon 1b (7), which is not associated with classical GOF mechanisms (i.e. DRPs and RNA foci). Additionally, two novel specific transcript variants (V4 and V5) have been reported in CD14+ myeloid cells, containing an could explain a preference of LOF over GOF alterations in microglia (69).

Astrocyte activation has been reported in *in vitro* models of C9orf72 HRE, with somewhat conflicting results. Our data show a general transcriptional silencing in astrocytes of both sALS and *C9orf72* HRE carriers, slightly more pronounced in the latter. This mostly involved transcriptional pathways related to synaptic and neuronal support, and is consistent with *in vitro* data from *C9orf72* iPSC derived astrocytes, showing a downregulation of gene networks involved in synaptic transmission and neuronal support, and increased expression of rRNA processing and ribosomal subunits (70). Contrary to our findings, in organoid systems, *C9orf72* astrocytes show a dysregulated transcriptional profile with increased levels of autophagy (71). alternative cryptic initial exon. These splice variants do not contain the HRE sequence and therefore

These inconsistencies may reflect intrinsic differences between human postmortem tissue, 2D and 3D models; but also the fact that microglia are missing in these cell culture platforms which may relevant aspects related to microglia-astrocytes crosstalk. Additional research using higher order organoids containing microglia, or xenotransplantation models will be useful to tackle these outstanding questions.

In summary, this study indicates that *C9orf72* HRE results in reduced *C9orf72* expression levels in the microglia and a subsequent reduction in their response to disease, consistent with deficits in phagocytic and lysosomal pathways. We hypothesize that this leads to a defective general glial response that extends to astrocytes, hindering a coordinated response that increases the risk for developing ALS.

## Acknowledgements

We are thankful to all participants and their relatives for their invaluable contributions to this research. We are indebted to the Department of Pathology at University Hospitals Leuven, The Netherlands Brain Bank (NBB) Amsterdam and HCB-IDIBAPS for sample and data procurement. We especially thank Mr. Michiel Kooreman, technical coordinator of NBB, for helping us with selecting and transferring the samples. We thank Mrs. Nikita Lamaire, ALS Clinical trial assistant, for helping with submission of this study protocol to the Ethics Committee Research UZ Leuven. We thank the VIB Nucleomics Core (www.nucleomics.be), specially Mr. Lim De Swert, Mrs. Kizi Coeck, Mrs. Jolien Vandewinkel and Mr. Stefaan Derveaux, for their help and expertise. P.M. has a research Fellowship of the European Academy of Neurology (no award/grant number). B.B. has a PhD fellowship of the University of Antwerp (DOCPRO4 FFB210307). R.M. receives funding from Fonds voor Wetenschappelijk Onderzoek (FWO grants no. G0C9219N, G056022N and G0K9422N), Alzheimer’s Association USA (2018-AARF-591110, 2018-AARF-591110-RAPID, ABA-22-968700). He also receives funding from Bright Focus Foundation (A2021034S), SAO-FRA (2021/0021), and the University of Antwerp (BOF-TOP 2022-2025). P.V.D. is a senior clinical investigator of the Flemish Fund for Scientific Research (T003519N) (FWO, Fonds voor Wetenschappelijk Onderzoek Flanders, Belgium) and is supported by the ALS Liga België and the KU Leuven funds “Een Hart voor ALS”, “Laeversfonds voor ALS Onderzoek” and “the Valéry Perrier Race against ALS fund (no award/grant number). This work was supported by C1 grants from KU Leuven (C14/17/107 and C14/22/132 to L.V.D.B, D.R.T, P.V.D), from FWO-Vlaanderen (G073222N and T003519N), the KU Leuven fund ‘Opening the Future’ and in part by the Intramural Research Programs of the NIH, National Institute on Aging (Z01-AG000949-02) and the National Institute of Neurological Disorders and Stroke. Computational resources and services used in this work were provided by the VSC (Flemish Supercomputer Center), funded by the Research Foundation - Flanders (FWO) and the Flemish Government. D.R.T receives additional funds from FWO (G0F8516N and G065721N) and SAO-FRA (2020/017).

## Authors contribution

P.M. designed all experiments and led this project; P.M., D.R. T and P.V.D collected the post-mortem human samples; D.R.T performed neuropathological analysis and dissected brain/spinal cord samples for analysis; P.M. performed RNA extraction with assistance of A.S.; S.K.P. performed the 10x genomics with the assistance of P.M., A.S. and N.H. B.B. and K.D performed the snRNAseq analysis; P.M., B.B., K.D, L.F., M.F., R.M. and P.V.D interpreted the snRNAseq data; P.M., B.B., R.M. and P.V.D. wrote the manuscript; L.F., L.V.D., M.F. and D.R.T critically revised the manuscript; all authors read and approved the final version of the paper.

## Tables

Supplementary Table 1. The demographic data of participants.

Supplementary Table 2. Overview of Microglia and Astrocytes gene markers.

Supplementary Table 3. Overview of DEG’s microglia in three conditions

Supplementary Table 4. Overview of DEG’s astrocytes in three conditions

## Methods (online)

### Human donor tissue

The brain and spinal cord samples from were obtained from the Department of Pathology at University Hospitals Leuven (UZ Leuven), the Netherlands Brain Bank (NBB), Netherlands Institute for Neuroscience (NIN), Amsterdam and Hospital Clinic de Barcelona (HCB) and the d’Investgiacions Biomédiques August Pi I Sunyer (IDIBAPS). All material has been collected from donors for or from whom a written informed consent for a brain autopsy and the use of the material and clinical information for research purposes had been obtained by the UZ Leuven (S65097, S59292, S60803) NBB and HCB-IDIBAPS.

### Experiments

#### RNA isolation and assessing of RNA integrity of post-mortem human brain and spinal cord tissues

1ml TriPure per 50-100 mg post-mortem tissue was added into a vial containing beads and post-mortem tissue. The sample was homogenized in tissue homogenizer (Magnalyser) 3 × 30 sec. at 6500 rpm. The homogenate was transferred to a new RNase free tube. 200 μl chloroform (= 1/5^th^ volume of TriPure) was added. The sample was mixed by inversion during 15 sec. and incubated at room temperature between 2 and 10 min. Next, the samples were centrifuged for 15 min. at 12000 rpm, 4°C. 3 phases became visible: aqueous phase (colorless) containing RNA, interphase (white color) containing DNA and organic phase (red color) containing protein. The aqueous phase was transferred to a new RNase free tube. Only in case of low expected yields of RNA, 1 μl glycogen was added and precipitated with 500 μl isopropanol (= 1/2th volume of TriPure). The sample was mixed by inversion (15 sec.), incubated at room temperature between 5 and 10 min., and centrifuged for 10 min. at 12000 rpm, 4°C. The supernatant was discarded. The pellet was washed with 1ml 75% EtOH (same volume as TriPure) and centrifuged for 5 min. at 12000 rpm, 4°C. Again, the supernatant was discarded. After drying of the pellet, RNase free water was added. Per sample, an amount of 1 μg of total RNA was used as input. RNA concentration and purity were determined spectrophotometrically using the NanoDrop ND-1000 (NanoDrop Technologies) and RNA integrity was assessed using a Bioanalyzer 2100 (Agilent).

#### Nuclei isolation of post-mortem human brain and spinal cord tissues

Recent advances in the isolation and sequencing of snRNA have demonstrated that, with optimal buffer/detergent compositions, the assay performance and gene detection sensitivity can be significantly enhanced. We used three different strategies including the protocol by standard nuclei isolation protocol Habib et al. (13), Tween with salts and Tris (TST) (14), and Soma-seq (15).

#### Standard nuclei isolation protocol

We used tissue pieces smaller than 50 mg for nuclei. They were transferred immediately in homogenizer containing 1 mL of ice-cold homogenization buffer (HB) (320 mM Sucrose, 5 mM CaCl_2_, 3 mM Mg(OAc)_2_, 10 mM Tris 7.8, 0.1 mM EDTA, 0.1% IGEPAL CA-360, 0.1 mM Phenylmethylsulfonyl fluoride, 1 mM β-mercaptoethanol with 5μl Rnasin Plus). The tissues were homogenized with 10 manual gentle strokes (pestle A) + 10 manual gentle strokes (pestle B). The homogenate was filtered through 70 μm cell mesh and was washed strainer with 1.65 ml HB and was added to a final volume of 2.65 ml. The nuclei was homogenated in the HB with 2.65 ml of Gradient Medium (5 mM CaCl_2_, 50% Optiprep, 3 mM Mg(OAc)_2_, 10 mM Tris 7.8, 0.1 mM Phenylmethylsulfonyl fluoride, 1 mM β-mercaptoethanol). 29% cushion was prepared by dilution of Optiprep with Optiprep Diluent Medium (150 mM KCl, 30 mM MgCl_2_, 60 mM Tris pH 8.0, 250 mM sucrose). The nuclei suspension in the HB + GM mix was layered over the 29 % cushion and centrifuged in an SW 41 Ti Swinging-Bucket Rotor (Beckman Coulter), at 7.700 rpm and 4°C for 30 min. The supernatant was removed with a Pasteur pipette andlower supernatant was removedwith P200. Nuclei was resuspended in 50 uL of Resuspension Buffer (PBS, 1% BSA) and transferred to a new tube. Resuspended nuclei was counted using LUNA dual florescence cell counter (Logos Biosystems).

#### TST nuclei isolation protocol

Similar to standard protocol above, tissue chunks less than 50 mg were used for nuclei isolation and transferred immediately in homogenizer containing 1 mL of ice-cold homogenization buffer (146 mM NaCl, 10 mM Tris 7.5, 1 mM CaCl_2_, 21 mM MgCl_2_, 0.03% Tween-20, 1% BSA, 25 mM KCl, 250 mM sucrose, 1 mM β-mercaptoethanol, 0.5X cOmplete™ Protease Inhibitor (Roche), 0.2U/ul RNasin® Plus Ribonuclease Inhibitor (Promega)). The tissues were homogenized with (n=15) manual gentle strokes with pestle A only. Nuclei suspension was filtered using 40 μm cell strainer. The filtrate was transferred to a low bind eppendorf and centrifuged at 500xg for 5 min. The supernatant was discarded. The pellet was resuspended thoroughly in 1 ml of HB w/o Tween buffer (146 mM NaCl, 10mM Tris 7.5, 1 mM CaCl_2_, 21 mM MgCl_2_, 1% BSA, 25 mM KCl, 250 mM sucrose, 1 mM β-mercaptoethanol, 0.5X cOmplete™ Protease Inhibitor (Roche), 0.2U/ul RNasin® Plus Ribonuclease Inhibitor (Promega), and transferred to 15 ml falcon. Additional 1.65 mL HB w/o tween buffer was added to a final volume of 2.65 ml. Rest of the protocol for density centrifugation and nuclei enrichment was identical to the standard protocol.

#### Soma-seq nuclei isolation protocol

The frozen tissue chunks were immediately transfered to homogenizer containing 1 mL of ice-cold homogenization buffer (10 mM Tris 8.0, 5 mM MgCl_2_, 25 mM KCl, 250 mM sucrose, 1 mM β-mercaptoethanol, 0.5X cOmplete™ Protease Inhibitor (Roche), 0.2U/ul RNasin® Plus Ribonuclease Inhibitor (Promega). The tissues were homogenized with (n=15) manual gentle strokes with pestle A only. The nuclei suspension was filtered using 70μm cell strainer. The the filtrate was transfered to a low bind eppendorf and centrifuged at 400xg for 5 min. The supernatant was discarded. RThe pellet was resuspended thoroughly in 1 ml of HB buffer, and transfered to 15 ml falcon. Additional 1.65mL HB buffer was added to a final volume of 2.65 ml. For this protocol as well the density centrifugation and nuclei enrichment is identical to what was done with standard nuclei isolation protocol.

#### Library preparation

Library preparations for the scRNA-seq was performed using 10X Genomics Chromium Single Cell 3’ Kit, v3.1 NextGEM chemistry (10X Genomics, Pleasanton, CA, USA). The cell count and the viability of the samples were assessed using LUNA dual fluorescence cell counter (Logos Biosystems) and a targeted cell recovery of 6000 cells were aimed for each of the samples. After cell count and quality control (QC), the samples were immediately loaded on to the Chromium Controller. Multiple samples (up to three patients) were pooled at equal counts and were loaded to a single lane of 10X. Single cell RNAseq libraries were prepared using manufacturers recommendations (Single cell 3’ reagent kits v3.1 user guide; CG000204 Rev D), and at the different check points the library quality was accessed using Qubit (ThermoFisher) and Bioanalyzer (Agilent). cDNA libraries were sequenced with an expected coverage of 50,000 reads per cell, either on Illumina’s NovaSeq 6000 platform using paired-end sequencing workflow and with recommended 10X; v3.1 read parameters (28-8-0-91 cycles).

## Data analysis

### Data pre-processing

Raw sequencing data was first demultiplexed using the cellranger (5.0.1) mkfastq command, the resulting fastq files were processed using the cellranger count command with the --include-introns option enabled and using the GRCh38-2020-A index provided on the 10X Genomics website. Cells passing the default cellranger filter were kept for subsequent analyses.

Samples that were multiplexed for droplet generation were demultiplexed into patient specific matrices using the following procedure. First, a library of SNPs present in the general population identified by the 1000 Genomes Project was obtained from http://ftp.1000genomes.ebi.ac.uk specifically, files ALL.chr(1-22,X,Y)_GRCh38_sites.20170504.vcf.gz were obtained and merged into a single vcf. Chromosomes were renamed to match the scheme used in the index provided by 10X genomics using bcftools annotate. These SNPs were then filtered to remove rare variants as well as to only include SNPs present in the regions of the genome included in the cellranger index using bcftools filter with the parameters --regions-file (with a bed file containing the above mentioned regions) and -i “INFO/AF[0] < 0.90 && INFO/AF[0] > 0.10”. This final vcf file was then used to run freemuxlet from the popscle package [https://github.com/statgen/popscle] with default parameters other than nsample = 2 and using the tools from https://github.com/aertslab/popscle_helper_tools to speed up computation. As freemuxlet only provides clusters of cells with no labels, we used verifyBamID [http://griffan.github.io/VerifyBamID/] to compare each genotype provided by freemuxlet to the bulk RNA-seq data unique to each patient, the highest scoring match was used to assign patient labels to freemuxlet clusters.

### Data processing and cell type annotation

Libraries were individually processed using Scanpy 1.7.2 (72). First, patient labels, and other associated metadata were added to the cells, with any doublets identified by freemuxlet being removed. Next, scrublet was used to remove any remaining identifiable doublets (threshold = 0.25) and all samples were combined into two main datasets based on their tissue of origin; motor cortex and spinal cord. Nuclei with less than 250 genes or more than 5% of total counts associated to mitochondrial genes were removed, the resulting matrices were normalised (to 10000 counts), log transformed, and highly variable genes were identified using default parameters. Both number of UMIs and percentage of mitochondrial reads were regressed out and the data was scaled to a maximum of 10. A principal component analysis (PCA) was used to reduce the dimensions of the data, Harmony (73) was used to integrate the PCA components based on the “Patient” batch key and the pcacv workflow from vsn-pipelines [https://doi.org/10.5281/zenodo.3703108] was then used to determine the number of principal components (PCs) to continue with (76 for Motor Cortex and 82 for Spinal Cord). Neighborhood embedding, UMAP and tSNE were calculated using the number of PCs listed above and clusters were identified at various resolutions (0.4, 0.8, 1.0, 1.2, 1.6, 2.0, 4.0, 8.0) using the leiden algorithm. The resulting data was exported to a loom file and uploaded to SCope (74), cell type annotations were provided via SCope by experts based on marker gene expression. All plots were generated with ggplot2 (v 3.3.5) in R v4.1.2.

### Processing of the microglia and astrocytes

To investigate microglia subsets, we isolated microglial nuclei from the single-nucleus data set (Masrori_Postmortem) for both the motor cortex and spinal cortex and merged them with Seurat (v4.1.0) under R version 4.1.2. The data was log normalized, highest variable genes were identified followed by scaling of the data and PCA. The data was batch corrected with Harmony (v0.1.0) based on patient identifiers and location. Cells were clustered using the first 20 principal components and UMAP plots were generated with Seurat standard methods. Non-microglial cells were excluded to perform final clustering of pure microglia.

When starting the reanalysis of the microglia extracted from the entire dataset, 5 other cell subsets were identified (peripheral macrophages n=898, synaptosome n= 821, oligodendrocytes n= 200 and proliferating cells n= 143). The clusters contained low expression of microglia markers CXCR3 and P2RY12 with subset specific markers for peripheral macrophages (*CD163, MRC1, F13A1*), oligodendrocytes (*MBP, SHROOM4, PLP1, PCDH9, EDIL3*) and synaptosomes/astrocytes (*SNAP25, RORA, DPP10, ROBO2, LSAMP*). The oligodendrocytes were most likely doublets of microglia and oligodendrocytes, which is why we still found overlap with the microglia markers. The same for the synaptosomes. Additionally, a cluster of proliferating cells (TOP2A, MIK67, MELK, KIF11) was excluded from the final analysis. While all the these subsets were initially defined as microglial cells in the large transcriptomic landscape of both the motor cortex and spinal cord, due to their transcriptomic similarity to microglia, the reanalysis of the microglial subset made these differences more apparent, driving us to exclude them.

Next, we extracted the astrocytes (n= 23483) from the spinal cord and motor cortex and re-identified the astrocyte population. Apart from astrocytes, two other clusters were identified: one of oligodendrocytes, n= 500 (*ELMO1, IL1RAPL1, SLC24A2, SHROOM4*) and one with doublets/low quality n =348 cells (*PCLO, WARS, RUNX1*), the latter being a cluster with markers of multiple cell types that could not be confidently annotated.

All plots were generated with ggplot2 (v 3.3.5) inR v4.1.2.

### Differential expression

Differential expression testing was performed using *DESEQ2* in Seurat Findmarkers. Genes with a p-value adjusted of <0.05 and log2FC of > +/− 0.20 were considered as differentially expressed. Heatmaps were generated with ComplexHeatmap_2.10.0.

### Gene enrichment analysis

Specific microglia functions were assessed by extracting human specific genes (taxon: 9606) 3 GO terms (GO:0005764, GO:0006954 and GO:006906) and 1 pathway in KEGG (M189). Gene signatures were calculated in Seurat with AddModuleScore.

### WGCNA

Matrices for microglia and astrocytes were extracted from each tissue and were used to perform WGCNA (75) to identify modules of co-expressed genes using the hdWGCNA package (v0.1.1.9005) (76). Genes were filtered to include only those expressed within at least 5% of the cells within a dataset using the SetupForWGCNA function, and MetacellsByGroups was used to create metacells by the combination of ALS Status, Patient and Library of origin using values of 25 for the number of cells to be combined and a maximum number of shared cells of 10, finally, metacells were normalized using NormalizeMetacells. Next, the groups of interest (Healthy, sALS and C9-ALS) were selected to build the co-expression network using SetDatExpr and TestSoftPowers was used per dataset to select a soft threshold (Spinal Cord – Astrocytes: 5, Spinal Cord – Microglia: 7, Motor Cortex – Astrocytes: 5, Motor Cortex – Microglia: 4). ConstructNetwork was used with default parameters to generate the co-expression networks and modules. Finally, module eigengenes were corrected for between-patient batch effects using harmony.

### WGCNA Modules-Trait Relationships

To identify WGCNA genes modules which may be correlated with ALS status, a one-way ANOVA (via the scipy python package v.1.7.3) was performed using the module eigengene values for each module across all datasets, p-values across all tests were corrected using the Benjamini/Hochberg correction (via the statsmodels python package v.0.12.2). A TukeyHSD test was then performed (also using statsmodels) on modules discovered in the ANOVA as having at least one significantly different mean with the goal of identifying the specific statuses which are different.

### Gene Ontology Enrichment Analysis

Gene ontology enrichment was performed using the g:OSt functionality of gProfiler (77) via the gprofiler-official python package (v1.0.0).

## Data availability

Processed data can be visualized via the interactive single cell data viewer SCope at https://scope.aertslab.org/#/Masrori_Postmortem.

Raw sequencing data have been deposited in the European Genome-phenome Archive (EGA) under and with data accession no. EGA… (to access the data itself under restricted access). Requests for accessing raw sequencing data will be reviewed by the VIB data access committee. Any data shared will be released via a Data Transfer Agreement that will include the necessary conditions to guarantee protection of personal data (according to European GDPR law).

Further information and requests for resources and reagents should be directed to and will be fulfilled by the Lead Contact, Philip Van Damme

## Supplementary figures

**Supplementary Figure 1.**
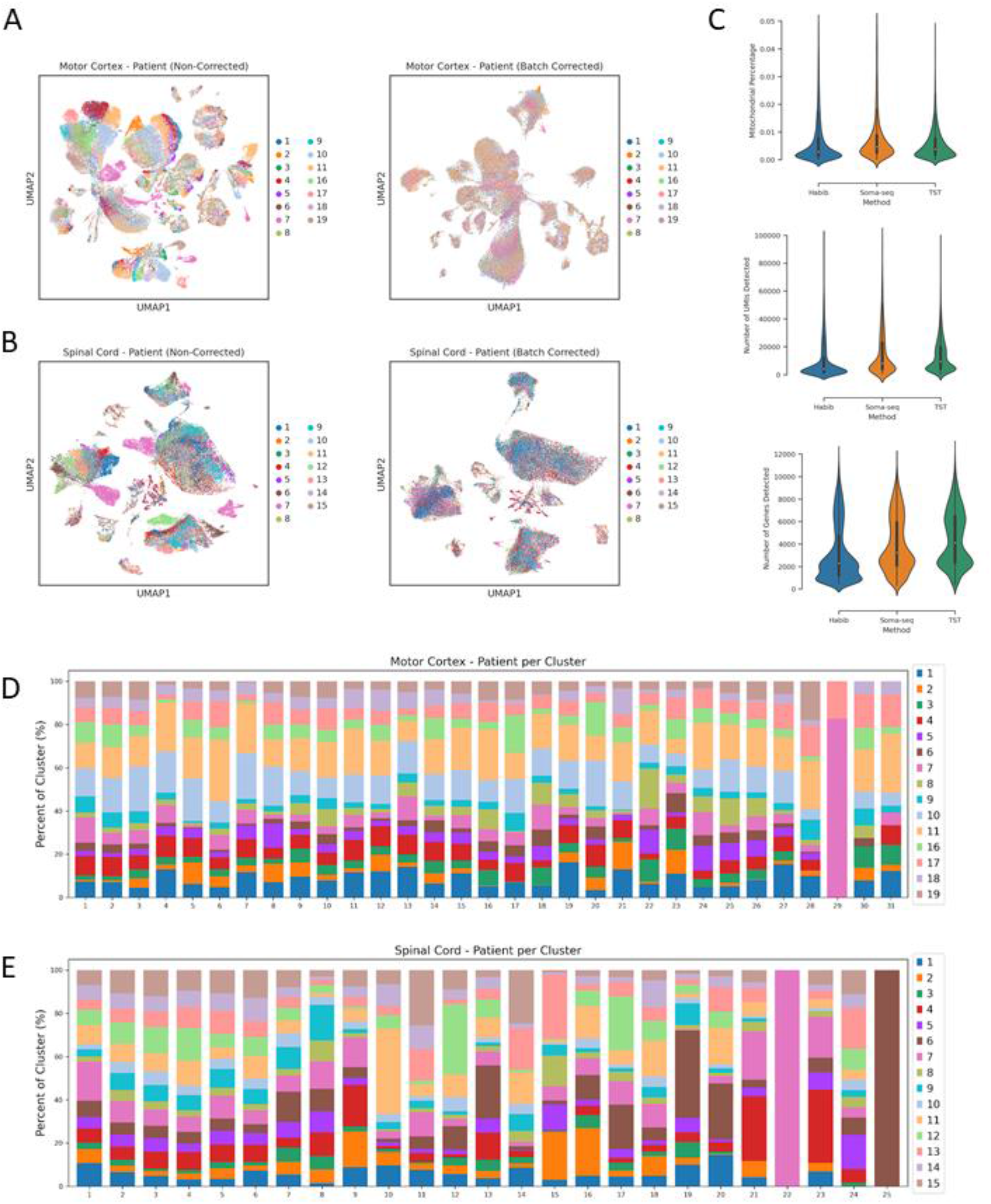
Overview of the full dataset and quality control. UMAP plot visualization of the single nuclei from (**a**) motor cortex and (**b**) prior and after batch correction. **(c)** Quality control of the three isolation protocols (Habib, Soma-seq, TST): percentage of mitochondrial RNA, number of molecules detected per cell, number of genes detected per cell. Contribution of each patient (x-axis) to each cluster of the (**d**) motor cortex and (**e**) spinal cord.

**Supplementary Figure 2:**
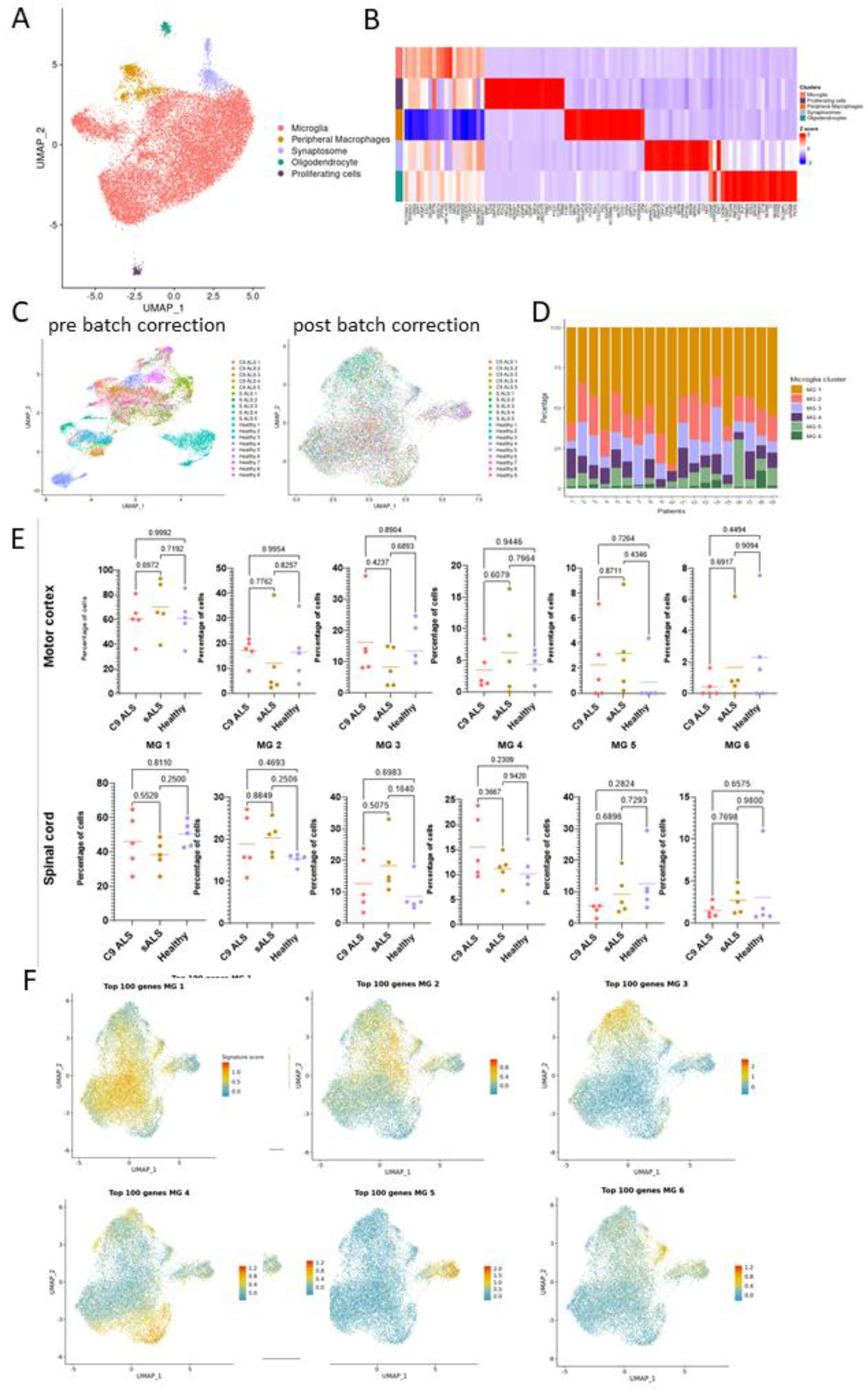
Extended clustering and quality control in the microglial subset. (**a**) UMAP plot visualizing 4,845 nuclei from the motor cortex 13,980 nuclei from the spinal cord. Cell populations are colored based on their assigned into the following clusters: microglia, peripheral macrophages, synaptosomes, oligodendrocytes and proliferating cells. (**b**) Heatmap showing the top 20 most expressed genes per cluster. (**c**) UMAP plot visualizing the selected microglia before and after batch correction. (**d**) Proportion of cells (y-axis) from each individual (x-axis) in the different microglia subsets. (**e**) proportion of cells (y-axis) from each genotype (per individual) in the different microglia subsets across both tissue locations. (f) Seurat module scores of the top 100 marker genes from the different microglia subsets.

**Supplementary Figure 3:**
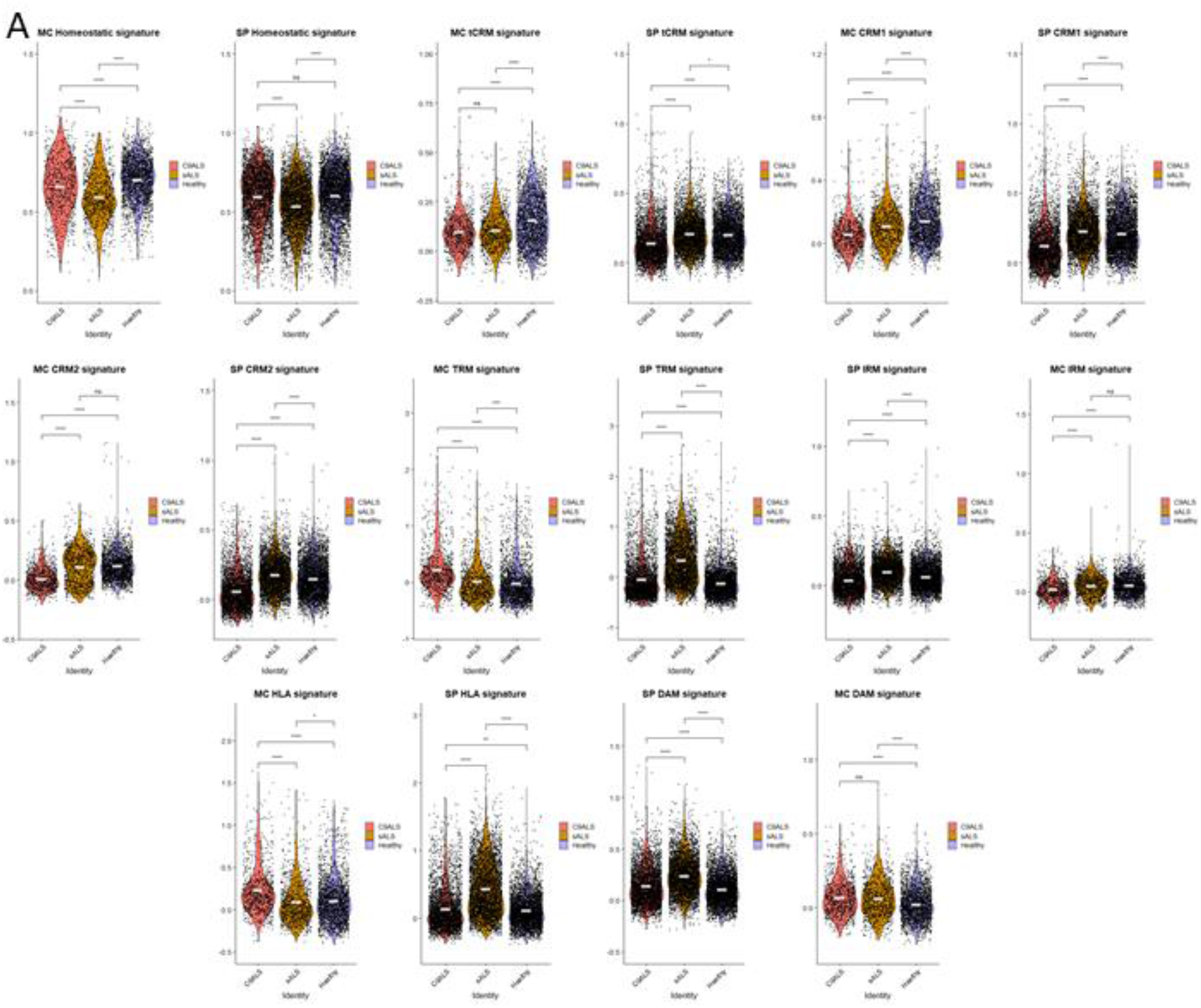
Extended analysis of the microglia transcriptional signatures. (**a**) Seurat module scores of homeostatic, transitioning cytokine response (tCRM), cytokine response 1 (CRM1), cytokine response 2 (CRM2), translational response (TRM), interferon response (IRM), antigen-presenting (HLA), disease-associated (DAM), and homeostatic microglia phenotypes as described by Mancuso et al., 2022.

**Supplementary Figure 4:**
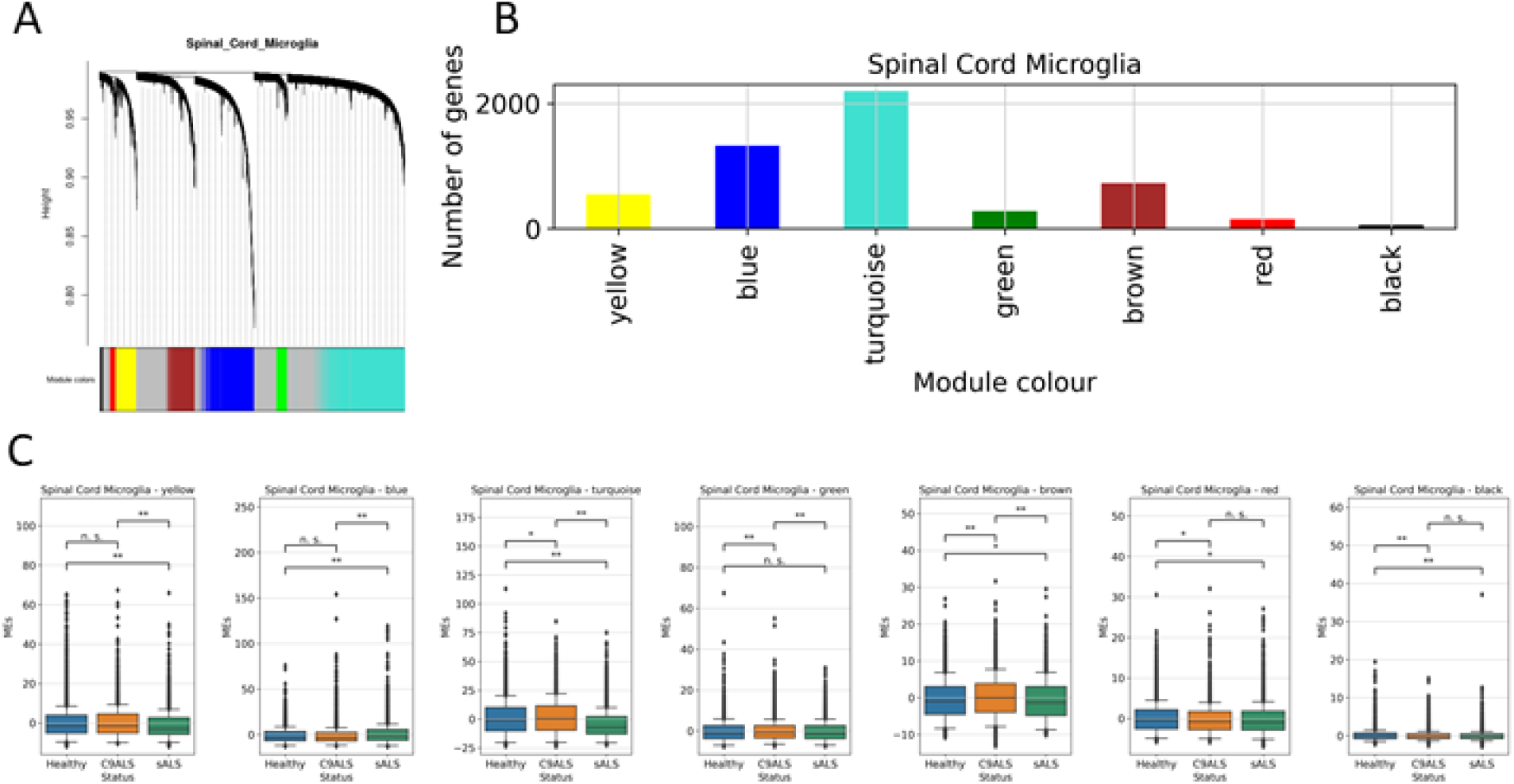
Extended WGCNA analysis. (**a**) Dendrogram showing result of unsupervised hierarchical clustering of modules identified by WGCNA. (**b**) Barplots depicting the number of genes per module. (**c**) Boxplot visualizing the module eigengenes (y-axis) per genotype (x-axis).

**Supplementary Figure 5:**
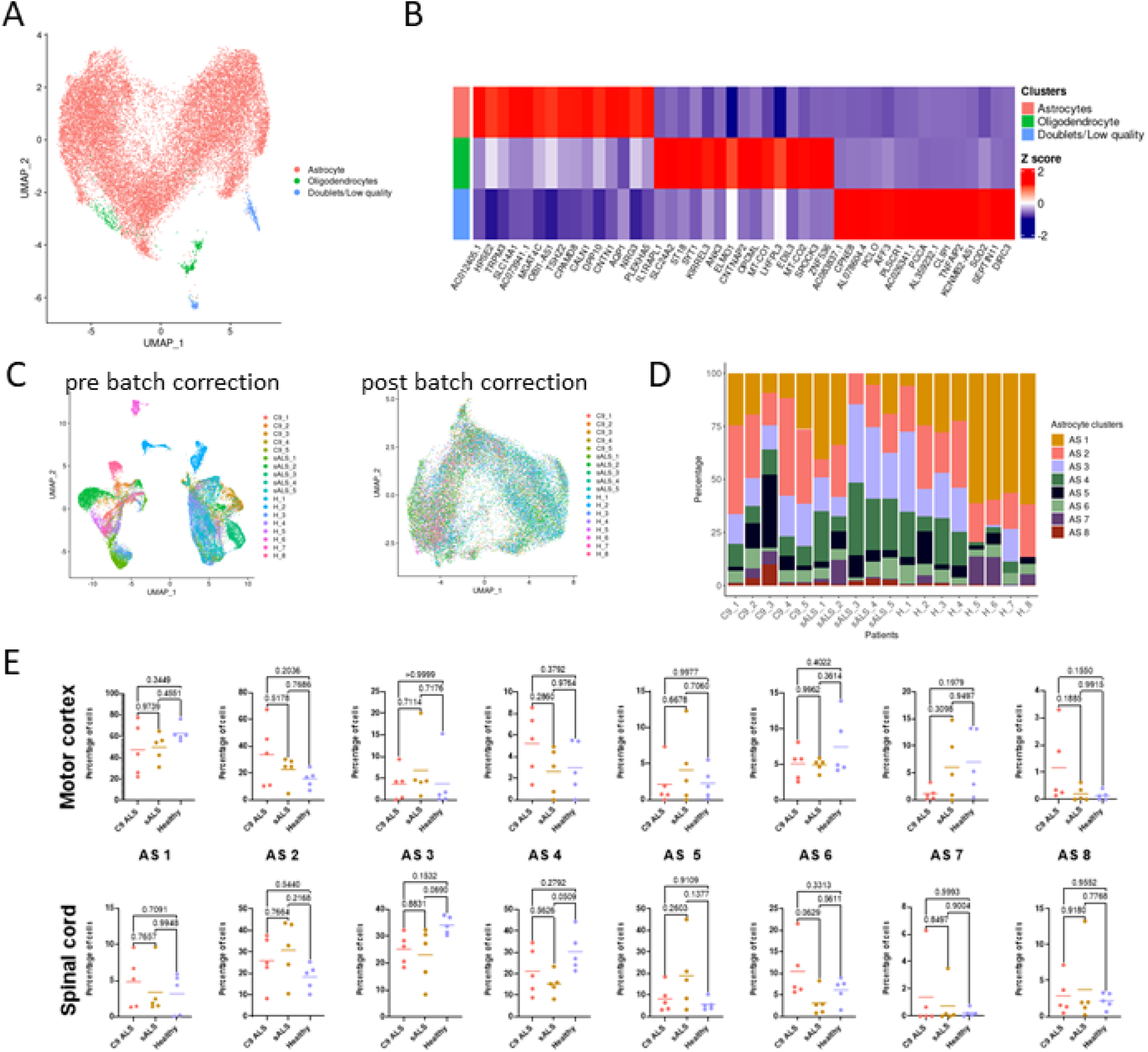
Extended clustering and quality control in the astrocytic subset. (**a**) UMAP plot visualizing 10,119 nuclei from the motor cortex 13,364 nuclei from the spinal cord. Cell populations are colored based on their assigned cluster into: astrocytes, oligodendrocytes, doublets/low quality. (**b**) heatmap showing the top 20 most expressed genes per cluster. (**c**) UMAP plot visualizing the selected astrocytes prior and after batch correction. (**d**) Proportion of cells (y-axis) from each individual (x-axis) contributing to the different astrocyte clusters. (**e**) Proportion of cells (y-axis) from each genotype (per individual) in the different astrocyte subsets across locations.

